# Dual Immune Checkpoint and Cytokine Receptor Modulation by an Engineered Human CTLA-4/IL-10 Bispecific Fusion Protein

**DOI:** 10.64898/2026.05.08.723457

**Authors:** Adi Cohen, Matan Gabay, Omri Cohen, Marina Sova, Amihai Lieberman, Anat Shemer, Nira Varda-Bloom, Elad Jacoby, Gal Cafri, Dorit Avni, Itamar Yadid, Maayan Gal

**Affiliations:** Department of Oral Biology, Goldschleger School of Dental Medicine, Gray Faculty of Medical & Health Sciences, Tel Aviv University, Tel Aviv, 6997801, Israel; Tel-Hai University, Upper Galilee, 1220800, Israel; Immunotherapy and genetic engineering lab, Sheba Medical Center, Tel-Hashomer, Israel; BMT Processing laboratory, Hematology Laboratory Sheba Medical Center, Tel-Hashomer, 5262100, Israel; Cell Therapy Lab, Sheba Medical Center, Tel-Hashomer, 5262100, Israel; Gray Faculty of Medical & Health Sciences, Tel Aviv University, Tel Aviv, 6997801, Israel; Division of Pediatric Hematology and Oncology, The Edmond and Lily Safra Children’s Hospital, Sheba Medical Center, Tel-Hashomer, 52621, Israel; Dina Recanati School of Medicine, Reichman University, Herzliya, 4610101, Israel; Migal - Galilee Research Institute, Kiryat Shmona, 11016, Israel

**Keywords:** Bispecific proteins, Protein engineering, Immunosuppression, CTLA-4, IL-10, Inclusion bodies, Virus-specific T cells (VSTs)

## Abstract

Bispecific fusion proteins represent a unique strategy for developing precision therapeutics. By linking functional domains from distinct proteins, these biomolecules can engage multiple targets, enhancing both therapeutic efficacy and safety. Unlike bispecific antibodies, low-molecular-weight fusion proteins offer distinct advantages, including reduced immunogenicity and superior tissue penetration due to their relatively compact size and structure. Such a profile is particularly valuable in managing complex inflammatory diseases, where modulating multiple pathways is required to impart an effective anti-inflammatory effect. Among the various regulators of immune signaling, the cytotoxic T-lymphocyte-associated protein 4 (CTLA-4) and interleukin-10 (IL-10) play imperative roles in immune suppression through their interactions with CD80/86 and IL-10R, respectively. While Fc-fused CTLA-4 is a clinically approved drug (e.g., Abatacept), the clinical development of IL-10 has been hampered by unpredictable immunostimulatory side effects. Here, we engineered a bispecific fusion protein linking the extracellular domain of CTLA-4 to IL-10. We successfully expressed the protein in *E. coli* as an N-terminal GST-tagged variant and refolded it from the inclusion bodies. Additionally, we achieved soluble expression of an Fc-tagged variant in mammalian CHO cells. Both origins demonstrated binding to their cognate receptors, CD80 and IL-10R. Finally, the fusion protein demonstrated a T cell-inhibitory effect by reducing Interferon-γ (IFNγ) secretion levels in an *in vitro* human Virus-Specific T cells (VSTs) model. This innovative protein engineering offers a promising strategy for addressing unmet clinical needs in autoimmune and inflammatory diseases.

## 1. Introduction

Bispecific biomolecules represent a paradigm shift in therapeutics, with the unique ability to bind two targets simultaneously [1, 2]. However, their design, expression, and evaluation remain challenging, often limited by poor stability and complex manufacturing requirements [3, 4]. Despite these hurdles, the clinical success of multiple bispecific agents, the majority of which are bispecific antibodies, has demonstrated enhanced specificity and reduced off-target effects compared with single-target therapeutics [5-9]. Thus, the engineering of bispecific biomolecules capable of modulating multiple pathways offers an attractive therapeutic strategy. This concept is particularly compelling in autoimmune diseases, which are driven by imbalances between immune-activating and immunosuppressive signaling cascades [10, 11].

The cytokine interleukin-10 (IL-10) and the checkpoint receptor cytotoxic T-lymphocyte-associated antigen-4 (CTLA-4) are two major regulators of immune homeostasis [12, 13]. IL-10 is a potent homodimer anti-inflammatory cytokine that signals through the heterodimeric IL-10 receptor (IL-10R), composed of the subunits IL-10RA and IL-10RB, to downregulate Th1 cytokine production, MHC class II dependent antigen presentation, and costimulatory molecules on macrophages [13, 14]. Owing to its broad activity across multiple immune cell types and its central role in balancing cytokine release, IL-10 is widely recognized as a key regulator of immune homeostasis in both health and disease [15]. These properties have motivated the clinical development of IL-10–based therapies for cancer and immune-mediated disorders [16-18]. However, clinical trials of IL-10 in inflammatory diseases such as Crohn’s disease have largely failed to show meaningful improvements in remission rates [19-21]. IL-10 limited therapeutic efficacy is often attributed to unanticipated immunostimulatory effects, which can promote prolonged B-cell survival, proliferation, and enhanced antibody production, thereby counteracting its intended anti-inflammatory purpose [22-25].

CTLA-4 and CD28 are well-characterized T-cell coreceptors belonging to the immunoglobulin (Ig) superfamily, each containing an Ig-like extracellular domain. Both receptors recognize the same ligands on antigen-presenting cells (APCs), CD80 (B7-1) and CD86 (B7-2) [26, 27]. However, they drive opposing signaling outcomes. CD28 engagement delivers a co-stimulatory signal required for T-cell activation, whereas CTLA-4 functions as a co-inhibitory checkpoint. By competing with CD28 for binding to CD80/CD86, CTLA-4 effectively dampens T-cell signaling and proliferation [28]. The reported binding affinity of CTLA-4 for CD80 (K_D_≈12 nM) is more than an order of magnitude stronger than that of CD28 (K_D_≈200 nM) [29]. Consequently, despite the high expression of CD28 on T cells, the higher affinity of CTLA-4 allows it to outcompete CD28 for ligand binding, leading to repression of T-cell proliferation, cell cycle progression, and cytokine production. Interestingly, both show lower affinity for CD86 [30, 31]. Exploiting this inhibitory activity, the extracellular domain (ECD) of CTLA-4 was engineered into an Fc-fusion protein, yielding clinically approved immunosuppressants such as Abatacept and Belatacept, which are now widely used to treat autoimmune diseases [32-34].

To address the clinical limitations of IL-10 while leveraging the proven efficacy of CTLA-4, we engineered a bispecific fusion protein that links the ECD of CTLA-4 to IL-10. This architecture is designed to enable simultaneous engagement of CD80 and IL-10R, effectively anchoring IL-10 to APCs and localizing its immunomodulatory activity. In this study, we describe the design, expression, and purification of the CTLA-4/IL-10 fusion protein in both bacterial (*E. coli*) and mammalian (CHO) systems, demonstrating that it retains functional, high-affinity binding to its cognate receptors. In addition, the protein efficiently reduced IFNγ secretion in *in vitro* Virus Specific T cells (VSTs) model.

## 2. Materials and methods

### 2.1 Gene synthesis and plasmid construction

DNA sequences encoding the ECD of human CTLA-4 (UniProt P16410) and IL-10 (UniProt P22301) were codon-optimized for *E. coli* expression or CHO cells and synthesized by GenScript. The fusion gene was constructed with a flexible glycine-serine linker separating the two domains and cloned into the pGEX-6P-1 vector with BamH1 and XhoI restriction enzymes. The final construct was designed to include an N-terminal GST tag and a C-terminal 6xHis tag. Full nucleotide and amino acid sequences are listed in Tables S1 and S2. For mammalian expression, the sequences were cloned into the pCDNA3 vector. These constructs were designed to feature an N-terminal secretion signal and a C-terminal Fc tag, facilitating secretion and stability in CHO cells. Full nucleotide and amino acid sequences for all constructs are listed in Tables S3 and S4.

### 2.2 Bacterial protein expression

Recombinant protein expression was performed in *E. coli* BL21 cells. Cultures were grown in LB medium supplemented with 100 μg/mL ampicillin at 37 °C with constant shaking at 200 RPM. Upon reaching an optical density (OD_600_) of 0.8, expression was induced by the addition of isopropyl β-D-1-thiogalactopyranoside (IPTG) to a final concentration of 1 mM. Following induction, cultures were incubated for 16 h at 25 °C with constant shaking at 200 RPM. Cells were harvested by centrifugation at 4,500 ×g for 45 min at 4 °C, and the resulting pellets were stored at -20 °C until further processing.

### 2.3 Inclusion body isolation, purification, and refolding

Purification of the bacterial fusion protein was performed at 4 °C unless otherwise noted. All buffer components are listed in Table S5. Cell pellets were resuspended in Lysis Buffer, sonicated (10 cycles of 30 s on/off at 40% amplitude using a Q125 sonicator, Qsonica), and centrifuged at 20,000 ×g for 20 min to harvest inclusion bodies. The pellet was solubilized in Denaturation Buffer with overnight stirring at room temperature. The solubilized fraction was centrifuged and purified using a 5 mL HisTrap FF column (Cytiva) on an ÄKTA go system. The column was equilibrated with Equilibration Buffer, washed with Wash Buffer, and the protein was eluted using Elution Buffer. Eluted fractions were dialyzed overnight against Refolding Buffer containing decreasing concentrations of urea (3 M and 0 M). Finally, the refolded protein was filtered (0.22 μm) and stored at -20 °C in Storage Buffer.

### 2.7 Mammalian cell culture and protein purification

Protein expression in CHO-S cells was executed by Genescript Ltd. Cells were maintained at 37 °C in a humidified atmosphere containing 8% CO_2_ on an orbital shaker. Plasmid transfection was performed by mixing DNA with the transfection reagent at an optimal ratio and adding the complex to the culture, followed by the addition of the media feed supplement 24 hours post-transfection. For purification, the cell mixture was loaded onto AmMag Protein A Magnetic Beads. The beads were washed twice with PBS (pH 7.2) and once with ddH2O. The protein was then eluted with 0.1 M glycine-HCl (pH 2.5–3.0), followed by immediate buffer exchange to a Sodium Acetate Buffer, 0.2 M L-arginine, pH 5.5.

### 2.4 SDS-PAGE and Western blot analysis

SDS–PAGE was utilized to assess the purity and molecular weight of the proteins. Samples were prepared by mixing with a loading buffer containing 0.2 M Tris-HCl, 8% (w/v) SDS, 6 mM bromophenol blue, 10% glycerol, and 1 mM β-mercaptoethanol (BME) as a reducing agent. For Western blot analysis, the purified protein was resolved on a 12% SDS-PAGE gel at 120 V for 80 min. Proteins were electroblotted onto a polyvinylidene difluoride (PVDF) membrane, which was subsequently blocked for 1 h at room temperature (RT) in PBS-T (PBS with 0.05% Tween 20) supplemented with 5% skim milk. The membrane was incubated overnight at 4 °C with the appropriate primary antibody (listed in Table S6) diluted in PBS-T. Following washing steps, the membrane was incubated for 1 h at RT with an HRP-conjugated secondary antibody (Table S6) in PBS-T. Immunoreactive bands were visualized using an Enhanced Chemiluminescence (ECL) substrate (Merck) and imaged via the ChemiDoc system (BioRad).

### 2.5 ELISA binding assay

ELISA was performed using a compatible PVC clear high bind ELISA microtiter 96-well plate (NEST). First, coating was made with GST-CTLA-4-IL-10-6xHis, diluted to 2 μg per well at a final volume of 100 μL. The plate was incubated ON at 4 °C, and afterward, it was washed 5 times with PBS-T 0.05%. 200 μL of blocking solution (5% skim milk in PBS-T 0.05%) was added and the plate was incubated for 5 h at RT to prevent non-specific binding. After incubation, the plate was washed five times with PBS-T 0.05%. Then, 0.2 μg of CD80 (His & AVI tag biotinylated, SinoBiological) or Fc-IL-10R (R&D Systems, USA) was added to the relevant wells at a final volume of 100 μL. Next, the plate was incubated at 4 °C ON. Following 5 washes with PBS-T 0.05%, blocking for 2 h, and additional washes, 100 μL of Fc-HRP-conjugated antibody or streptavidin-HRP (1 μL of antibody diluted in 7 mL PBS-T) was added to the relevant wells at a final volume of 100 μL and incubated at room temperature for 1 hour. Washing steps were repeated, then 100 μL of TMB (SurModics) were added to each well and the plate was incubated at RT up to 1 h. After color development, 50 μL of 0.5 M H_2_SO_4_ was added to each well to stop the enzymatic reaction. Plate absorbance was immediately measured using a plate reader (BioTek Synergy H1) at an absorbance of 450 nm.

### 2.6 Biolayer interferometry (BLI) affinity measurements using Octet96

Affinity measurements of CTLA-4-IL-10 protein to CD80 and IL-10R were performed using Octet96 (Satorius Ltd). Measurements were at 25 °C in PBS supplemented with 0.02% Tween-20 and 0.1% BSA to minimize non-specific binding. For binding measurements, 5 ng/mL of biotinylated human CD80 (BPS, Cat# 71114) or 2.5 ng/mL of IL-10R (Antibodies online, Cat# ABIN7504430) were immobilized onto streptavidin biosensors. Serial dilutions of the protein (ranging from 0.01 µM to 2 µM) were prepared in the same buffer and loaded for 500 s to record the association phase, followed by dissociation for an additional 600 s in buffer alone.

### 2.7 Generation of virus-specific T-cells

The method was adapted from previously published protocols [35-38]. Briefly, fresh peripheral blood mononuclear cells (PBMCs) from a healthy donor were isolated from whole blood by density-gradient centrifugation using Lymphoprep™ Ficoll (Serumwerk Bemburg AG). PBMCs were seeded in 24-well plates (3 × 10^6 cells per well) (Greiner) in virus-specific T cell (VST) medium consisting of a 1:1 mixture of AIM V (Gibco) and RPMI 1640 (Sartorius), supplemented with 1% L-glutamine (Sartorius), 1% penicillin–streptomycin (Sartorius), and 10% human serum (Access Biologicals). Cells were stimulated in vitro with an adenovirus Hexon peptide library (Miltenyi Biotec) at a final concentration of 1 µg/mL for 2 hours, followed by incubation at 37 °C in 5% CO_2_. After 2 hours, fresh medium was added, and cells were cultured for an additional 3 days under the same conditions. Every 3 days thereafter, cells were expanded into larger culture vessels with fresh VST medium supplemented with interleukin-2 (IL-2; 300 IU/mL; Clinigen) and maintained at 37 °C in 5% CO_2_. Cells were expanded for 10–12 days for VST enrichment before cryopreservation.

### 2.8 Co-culture assay for quantifying IFNγ levels

PBMCs from the same donor were thawed and seeded at 7 × 10^5 cells per well in 96-well plates (Greiner) containing VST medium. Cells were incubated for 2 hours at 37 °C with 5% CO_2_ to allow adherence. Non-adherent cells were removed by washing, leaving a population enriched by antigen-presenting cells (APCs; ±10% of total cells). APCs were loaded with the Hexon peptide library (Miltenyi Biotec) at either 200 ng/mL or 1 ng/mL for 2 hours at 37 °C with 5% CO_2_. Thirty minutes before co-culture, VSTs were seeded at 1×10^5 cells per well in a separate 96-well plate with VST medium. The plate was centrifuged at 1,300 rpm for 5 minutes; the supernatant was removed, and VSTs were resuspended in medium containing IL-10, CTLA-4, or CTLA-4/IL-10 at 200 nM, then incubated together for 30 minutes at 37 °C. Following antigen loading, APCs were washed with PBS, and the VSTs containing proteins were added to the APC-containing wells to initiate co-culture. Cells were incubated for 18 hours at 37 °C with 5% CO_2_. After incubation, culture supernatants were collected and analyzed for interferon-γ (IFN-γ) secretion by ELISA according to the manufacturer’s protocol (BioLegend).

## 3. Results

### 3.1 Evaluating the bacterial expression of GST-tagged bispecific CTLA-4-IL-10

In our first step, we evaluated the expression and purification of the bispecific fusion protein in *E. coli*, chosen for its cost-effectiveness and rapid scalability as a protein production host [39]. Although *E. coli* offers a fast, inexpensive, and robust expression system, it lacks eukaryotic post-translational modifications and frequently yields recombinant proteins in inclusion bodies. To mitigate potential folding issues and facilitate purification, we designed the construct with an N-terminal glutathione S-transferase (GST) tag and a C-terminal hexahistidine (His×6) tag [40, 41]. The gene encoding the bispecific protein was cloned into the pGEX-6P-1 vector and transformed into *E. coli* BL21 cells. The resulting ∼60 kDa fusion protein comprised, from N-to C-terminus: GST, an HRV-3C protease cleavage site, the CTLA-4 extracellular domain (ECD), a flexible linker, IL-10, and a 6×His tag (Fig. 1A; Supplementary Tables S1 and S2). To assess expression, cultures were grown to an OD_600_ of 0.8 and induced with 1 mM IPTG under three conditions: 37 °C for 4 h, or 25 °C and 15 °C overnight. Expression was evaluated by SDS–PAGE analysis of soluble and insoluble fractions. Under all tested conditions, only a faint band was observed in the soluble fraction, whereas the majority of the recombinant protein was detected in the insoluble pellet, indicating predominant localization to bacterial inclusion bodies (Fig. 1B). Schematic representations of the multi-component fusion protein with and without the GST tag are shown in Figures 1C and 1D, respectively.

**Figure 1.**
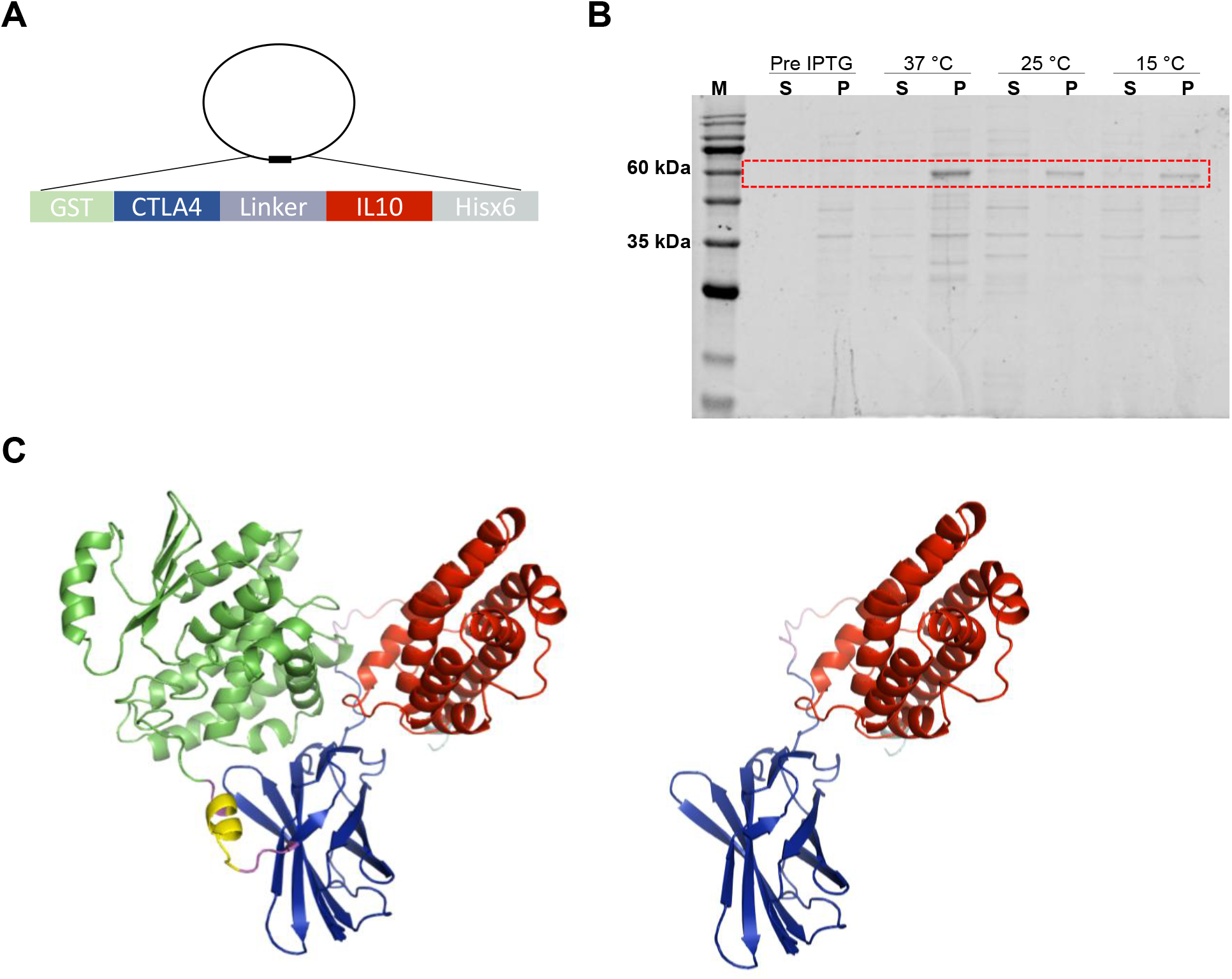
Construct design, bacterial expression analysis and structural modeling of the bacterial CTLA-4/IL-10 bispecific protein. (**A)** Schematic representation of the bacterial expression plasmid encoding the bispecific fusion protein comprising an N-terminal GST tag, the CTLA-4 extracellular domain, a flexible linker, IL-10, and a C-terminal Hisx6 tag. **(B)** SDS–PAGE analysis of protein expression following induction at 37 °C, 25 °C, or 15 °C. Lane M, molecular weight marker; “S” and “P”, indicate the soluble and insoluble (pellet) fractions, respectively. The red dashed rectangle marks the band at the expected molecular weight (∼60 kDa). (**C)** AlphaFold-predicted models of CTLA-4–IL-10 with and without the N-terminal GST tag. Domains are color-coded as follows: CTLA-4 (blue), IL-10 (red), GST (green), Hisx6 (cyan), HRV-3C cleavage site (yellow), and linker (magenta).

### 3.2 Bacterial CTLA-4/IL-10 purification and refolding

To isolate the bispecific fusion protein, inclusion bodies harvested from a 1 L culture were solubilized in denaturing buffer containing 6 M guanidine hydrochloride. After clarification of the lysate by centrifugation, the denatured lysate was subjected to Ni–NTA affinity chromatography under denaturing conditions (Fig. 2A). Although a fraction of the target protein was detected in the flow-through, a distinct purified band at the expected molecular weight was enriched and eluted in the elution fraction (Fig. 2A, lane 4). To restore native folding, the eluted protein was refolded by stepwise dialysis against buffers containing decreasing urea concentrations (3 M to 0 M). Arginine and a cystine/cysteine redox pair were included to maintain solubility and promote correct disulfide bond formation. The refolded protein was subsequently filtered (0.22 μm), and SDS–PAGE analysis confirmed recovery of the CTLA-4/IL-10 fusion protein (Fig. 2B).

**Figure 2.**
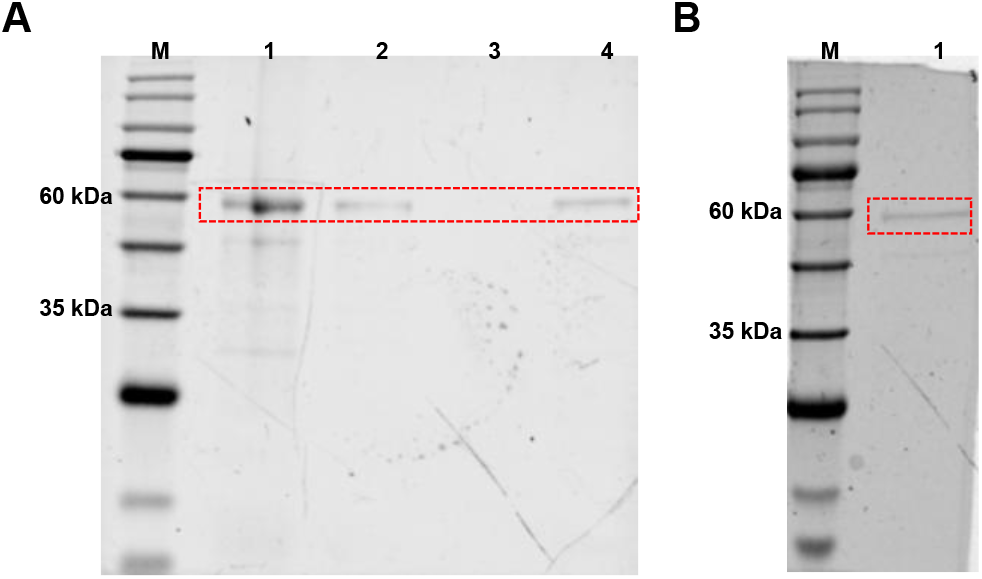
Purification and refolding of bacterially expressed CTLA-4/IL-10. **(A)** SDS–PAGE analysis of Ni–NTA affinity purification performed under denaturing conditions. Lane M, molecular weight marker; lanes 1–4, load, flow-through (FT), wash, and elution fractions, respectively. **(B)** SDS–PAGE analysis of the eluted purified CTLA-4/IL-10 fusion protein after refolding. The red dashed rectangle marks the band corresponding to the CTLA-4/IL-10 protein. Lane M, molecular weight marker; lane 1, soluble fraction following refolding.

To assess the structural integrity of the refolded fusion protein and confirm expression of its individual domains, we performed Western blotting using antibodies against the 6×His, GST, CTLA-4, and IL-10 (Fig. 3). All four antibodies detected a band at the expected molecular weight, consistent with the intact CTLA-4/IL-10 fusion protein and confirming the presence of each component. Notably, a lower–molecular-weight species was also detected by SDS–PAGE and Western blotting. This band may reflect partial proteolysis of the fusion protein, potentially arising from nonspecific cleavage within the N-terminal region or at the C-terminus.

**Figure 3.**
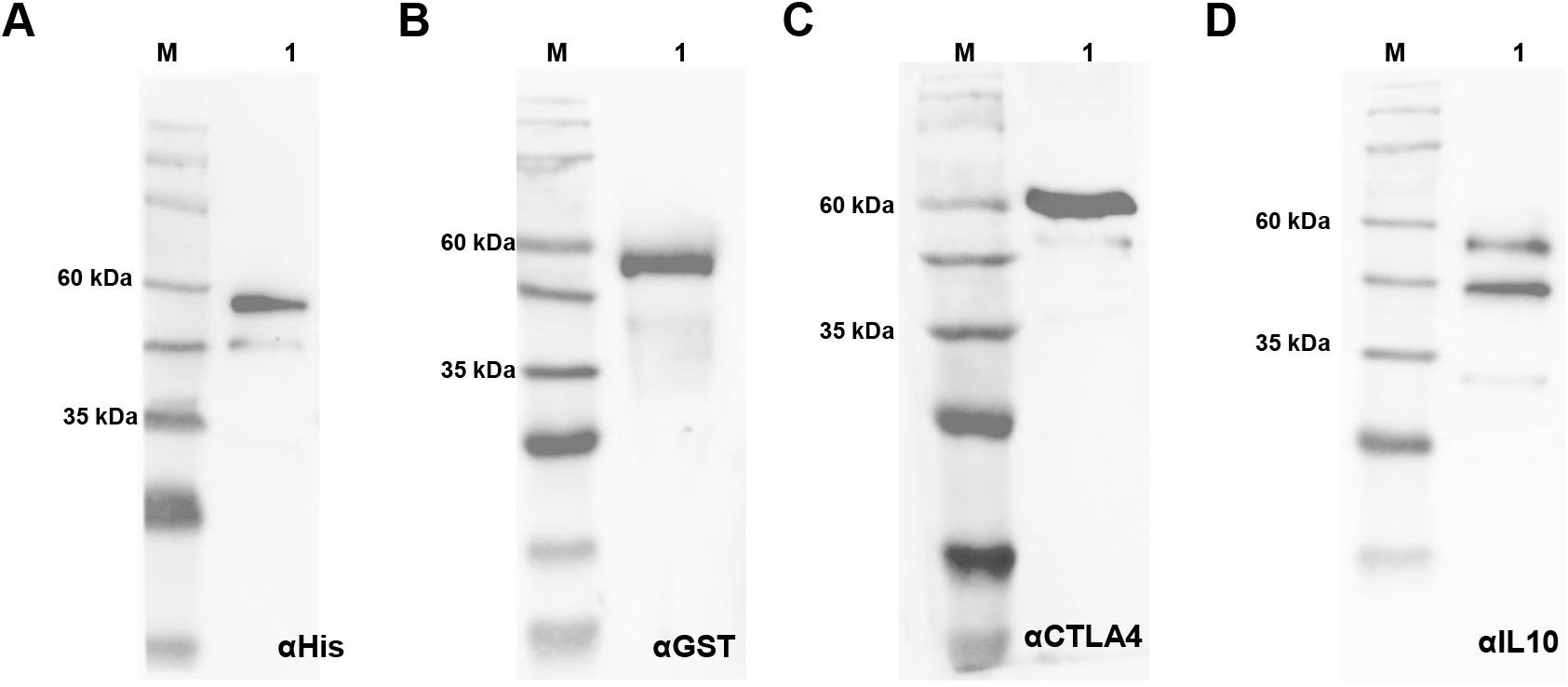
Western blot analysis of the refolded CTLA-4/IL-10 bacterial protein. Immunoblotting was performed to confirm the presence of the major domains within the refolded fusion protein. Membranes were probed with antibodies against **(A)** the 6×His, **(B)** GST, **(C)** CTLA-4, and **(D)** IL-10. The predominant band at ∼60 kDa corresponds to the full-length CTLA-4–IL-10 fusion protein. Lane M, molecular weight marker; lane 1, refolded CTLA-4/IL-10.

### 3.3 Binding of bacterial CTLA-4/IL-10 to CD80 and IL-10R

To evaluate the dual-binding capability of the refolded fusion protein, we performed an ELISA using its cognate receptors, CD80 and IL-10R (Fig. 4A). The GST-tagged bispecific protein was immobilized on 96-well plates (2 μg/well) overnight. After washing and blocking, wells were incubated with either biotinylated CD80 or Fc-tagged IL-10R (0.2 μg/well). Binding was detected using HRP-conjugated streptavidin (for CD80) or an HRP-conjugated anti-Fc antibody (for IL-10R). Signals were developed with TMB substrate and quantified at 450 nm following quenching with 0.5 M H_2_SO_4_. As shown in Figures 4B and 4C, the bispecific protein (black bars) exhibited robust binding to both CD80 and IL-10R, whereas the GST-only control (white bars) produced background-level signal. Together, these results indicate that the CTLA-4 and IL-10 domains remain functional after refolding, supporting the specific recognition of both receptors.

**Figure 4.**
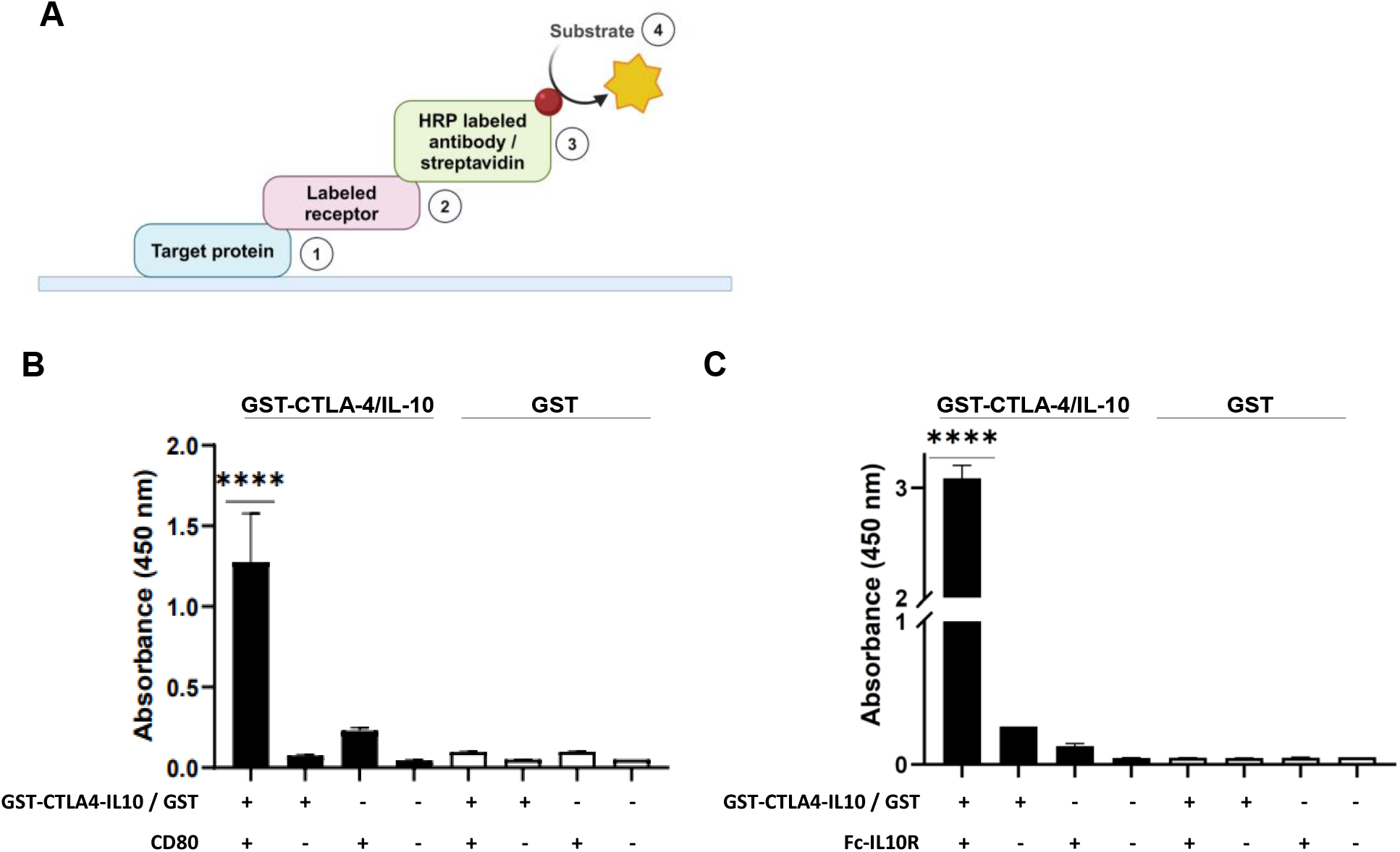
ELISA-based analysis of bacterial CTLA-4/IL-10 binding to CD80 and IL-10R. **(A)** Schematic illustration of the experimental workflow (created with BioRender.com). **(B-C)** Specific binding of the GST-tagged bispecific protein (black bars) compared with GST alone as a control (white bars) to **(B)** CD80 and **(C)** Fc-IL-10R. Data are presented as mean ± SD (n = 3). Statistical significance was determined by one-way ANOVA using GraphPad Prism (****, p < 0.0001).

### 3.4 Binding affinity of bacterially produced CTLA-4/IL-10 to CD80 and IL-10R measured by biolayer interferometry

To quantify the binding affinity of the bacterially produced CTLA-4/IL-10 fusion protein to its cognate receptors, we performed biolayer interferometry (BLI) using an Octet RED96 system (Fig. 5). Streptavidin biosensors were loaded with the biotinylated IL-10R (Fig. 5A) or CD80 (Fig. 5B) and kinetic interactions were measured by exposing the biosensors to the refolded protein CTLA-4/IL-10 as analyte at concentrations ranging from 0.01 to 2 μM. GST alone was included as a negative-control analyte to assess nonspecific interactions. Association and dissociation were monitored for 500 s and 600 s, respectively, enabling estimation of K_on_ and K_off_ values (Table 1). Sensorgrams were globally fitted to a 1:1 binding model. The fusion protein bound specifically to both targets, yielding equilibrium dissociation constants (K_D_) of 155.3 nM for CD80 and 45.7 nM for IL-10R.

**Table 1.** Kinetic binding parameters for bacterially produced CTLA-4–IL-10 interacting with immobilized CD80 or IL-10R measured by BLI. Values are mean ± SD from three independent experiments. *K*_*D*_ *calculated as K*_*off*_*/K*_*on*_

| Bacterial CTLA4/IL10 | Kd (nM) | Kon (1/Ms) | Koff (1/s) |
| --- | --- | --- | --- |
| Binding to CD80 | 155.33 (±13.65) | 2.035E03 (±4.79E02) | 3.523E-04 (±6.21E-05) |
| Binding to IL10R | 45.7 (±2.44) | 3.382E03 (±1.2E02) | 1.548E-04 (±1.467E-05) |

**Figure 5.**
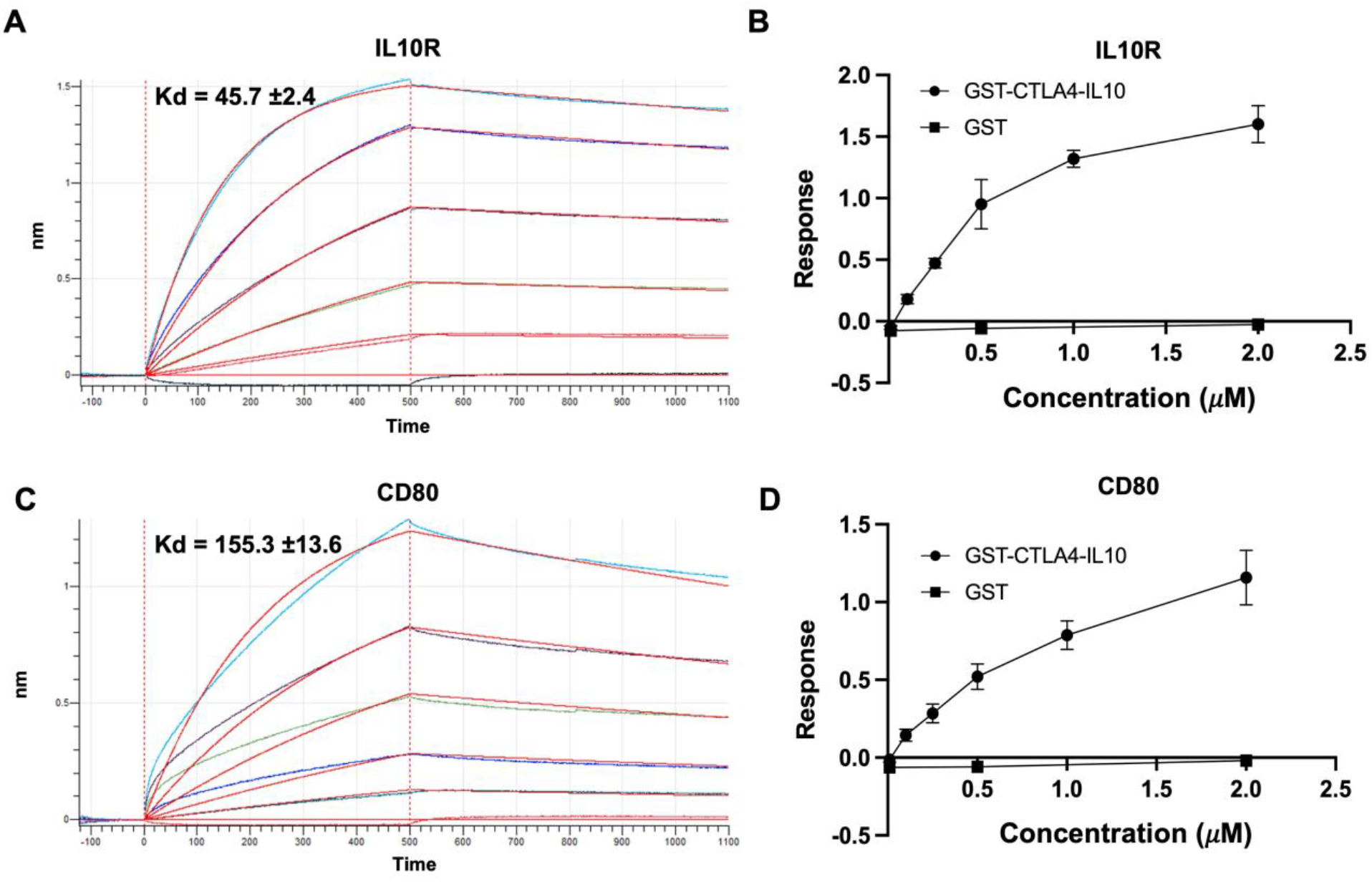
Kinetic binding analysis of CTLA-4/IL-10 by BLI. **(A)** Representative sensorgrams and kinetic fits, and **(B)** steady-state binding analysis for CTLA-4–IL-10 binding to immobilized IL-10R. **(C)** Representative sensorgrams and kinetic fits, and **(D)** steady-state binding analysis for CTLA-4–IL-10 binding to immobilized CD80. Biotinylated receptors were loaded onto streptavidin biosensors and incubated with CTLA-4/IL-10 at concentrations ranging from 0.01 to 2 μM in PBS + 0.02% Tween-20. Association and dissociation were recorded for 500 s and 600 s, respectively. GST alone was tested as a negative-control analyte under the same conditions. Data were globally fitted to a 1:1 binding model. Values represent mean ± SD from three independent experiments.

### 3.5 Expression and purification of Fc-tagged CTLA-4/IL-10 fusion proteins in mammalian cells

The experiments with the bacterial protein confirmed its functionality which establishes a potential robust system to explore the effect of genetic variations. However, while bacterial expression enables rapid screening, mammalian expression is typically required for in-vivo assays and therapeutic development because it supports proper disulfide bond formation and native glycosylation, which can strongly influence protein stability, pharmacokinetics, immunogenicity and receptor interactions. Accordingly, we expressed three Fc-tagged CTLA-4/IL-10 fusion variants in CHO-S cells (Fig. 1A). The constructs with an N-terminus secretion signal were i) CTLA-4–IL-10–Fc, ii) IL-10–CTLA-4–Fc, and a dual-cytokine variant in which two IL-10 domains were fused in tandem via a short linker iii) IL-10–IL-10–CTLA-4–Fc. SDS–PAGE under reducing conditions confirmed that all three variants were efficiently secreted and migrated at their expected molecular weights (Fig. 6B–D). In contrast, non-reducing SDS–PAGE revealed marked differences in oligomeric assembly. Both CTLA-4–IL-10–Fc (Fig. 6A) and IL-10–CTLA-4– Fc (Fig. 6B) exhibited multiple high–molecular-weight species, consistent with heterogeneous oligomerization and/or aggregation. By comparison, IL-10–IL-10–CTLA-4–Fc migrated predominantly as a single, well-defined band at the expected homodimer size (∼160 kDa), indicating improved assembly and minimal aggregation (Fig. 6C). These data suggest that the forced dimerization of the IL10 domain strongly influences folding and solubility and promotes a more homogeneous Fc-fusion product. Structural models of the tandem IL-10 construct are shown with the C-terminal Fc domain (Fig. 6E) and without Fc (Fig. 6F).

**Figure 6.**
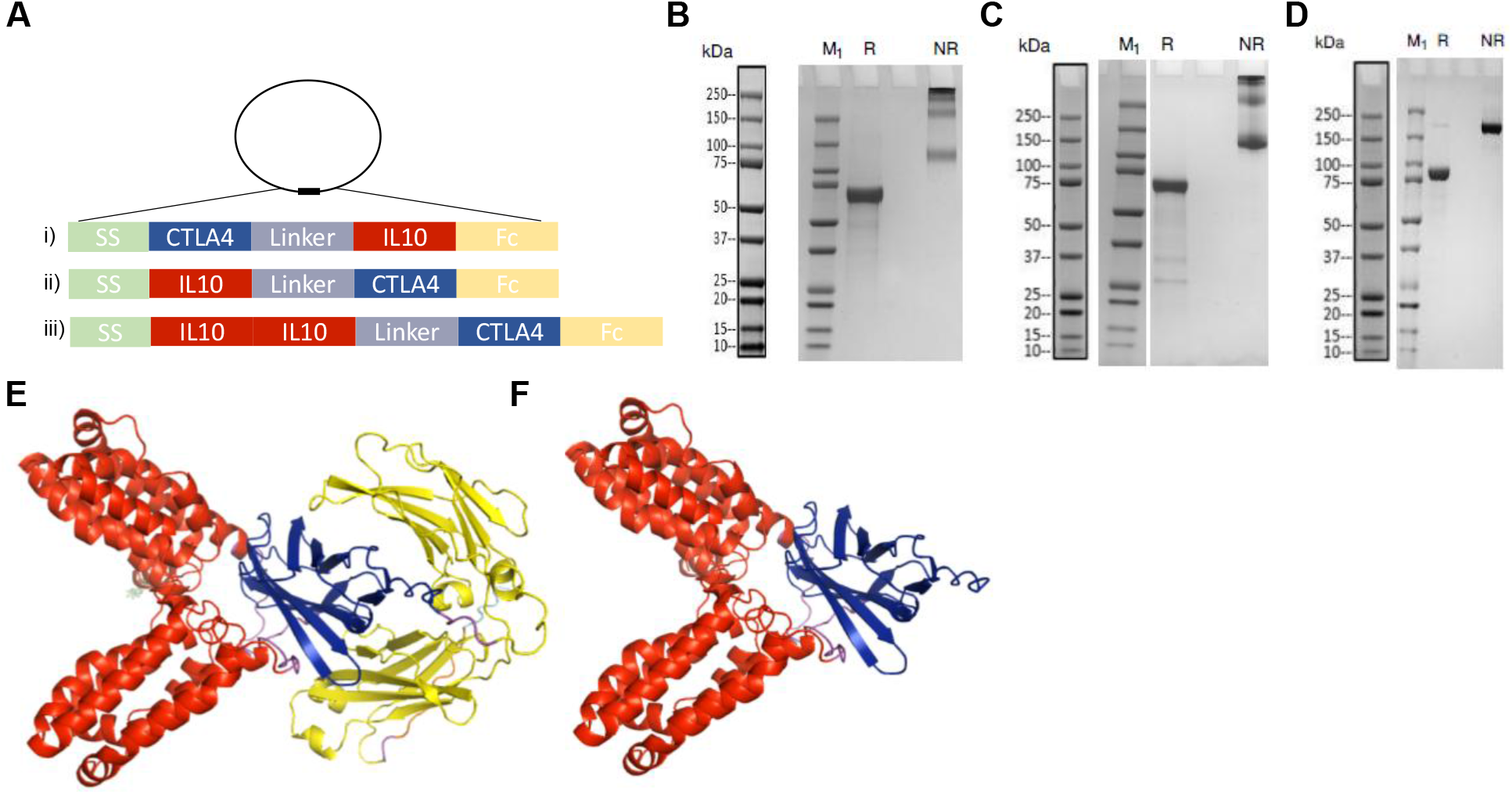
SDS-PAGE analysis of CTLA-4/IL-10 variants expressed in CHO-S cells. All fusion proteins were engineered with an N-terminal secretion signal and a C-terminal Fc tag. Panels show the migration profiles of **(A)** CTLA-4-IL-10, **(B)** IL-10-CTLA-4, and **(C)** the tandem variant IL-10-IL-10-CTLA-4. Lanes: M, molecular weight marker; R, reduced conditions; NR, non-reduced conditions. (D) and (E) Alpha Fold model of the IL-10-IL10-CTLA-4 structure with and without the Fc domain. Color code: Red – IL-10; Blue – CTLA-4; Yellow – Fc.

### 3.6 Binding of the IL10-Il10/CTLA-4 produced in mammalian cells to CD80 and IL-10R

Based on the superior oligomeric stability of the dual IL-10/CTLA-4 variant, we selected this construct for functional evaluation via ELISA. The fusion protein was immobilized on 96-well plates at a concentration of 2 μg/well. To avoid cross-reactivity with the construct’s Fc domain, binding was assessed using 0.2 μg/well biotinylated CD80 (**Fig. 7A**) or biotinylated IL-10R (**Fig. 7B**). Interactions were subsequently detected using HRP-conjugated streptavidin. Signals were developed with TMB substrate and quantified at 450 nm after acid quenching.

**Figure 7.**
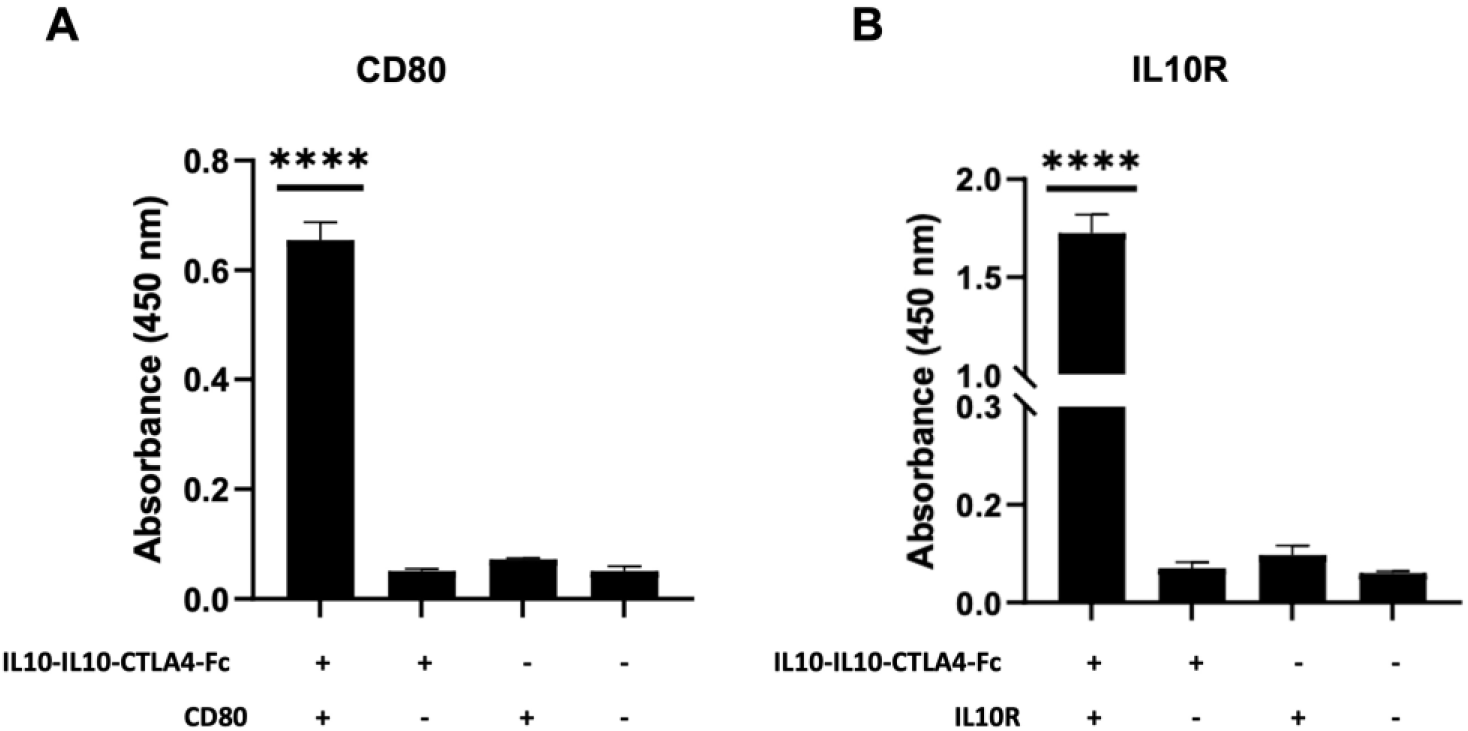
ELISA-based binding of the IL-10–IL-10/CTLA-4–Fc variant produced in mammalian cells to CD80 and IL-10R. The Fc-fusion protein was immobilized on 96-well plates and incubated with biotinylated (A) CD80 or (B) IL-10R. Binding was detected using HRP-conjugated streptavidin. Data are presented as mean ± SD. Statistical significance was determined by one-way ANOVA using GraphPad Prism (****, p < 0.0001).

### 3.7 Binding affinity of the IL-10–IL-10/CTLA-4–Fc produced in mammalian cells to its receptors via Biolayer Interferometry

To quantify the binding kinetics and steady-state affinity of the IL-10–IL-10/CTLA-4–Fc fusion protein produced mammalian in cells for its cognate receptors, we performed BLI using the same experimental conditions as for the bacterial protein (Fig. 8). Biotinylated receptors were immobilized on streptavidin biosensors and incubated with the mammalian cells produced fusion protein at concentrations ranging from 0.01 to 2 μM in PBS supplemented with 0.02% Tween-20. Association and dissociation were monitored for 500 s and 600 s, respectively, enabling estimation of k_on_ and k_off_ values (Table 2), and sensorgrams were globally fitted to a 1:1 binding model. The mammalian cells produced a fusion protein bound to CD80 with an apparent equilibrium dissociation constant (K_D_) of 462.8 ± 11.98 nM, and exhibited extremely tight binding to IL-10R with a K_D_ below the detection limit (<1 pM).

**Figure 8.**
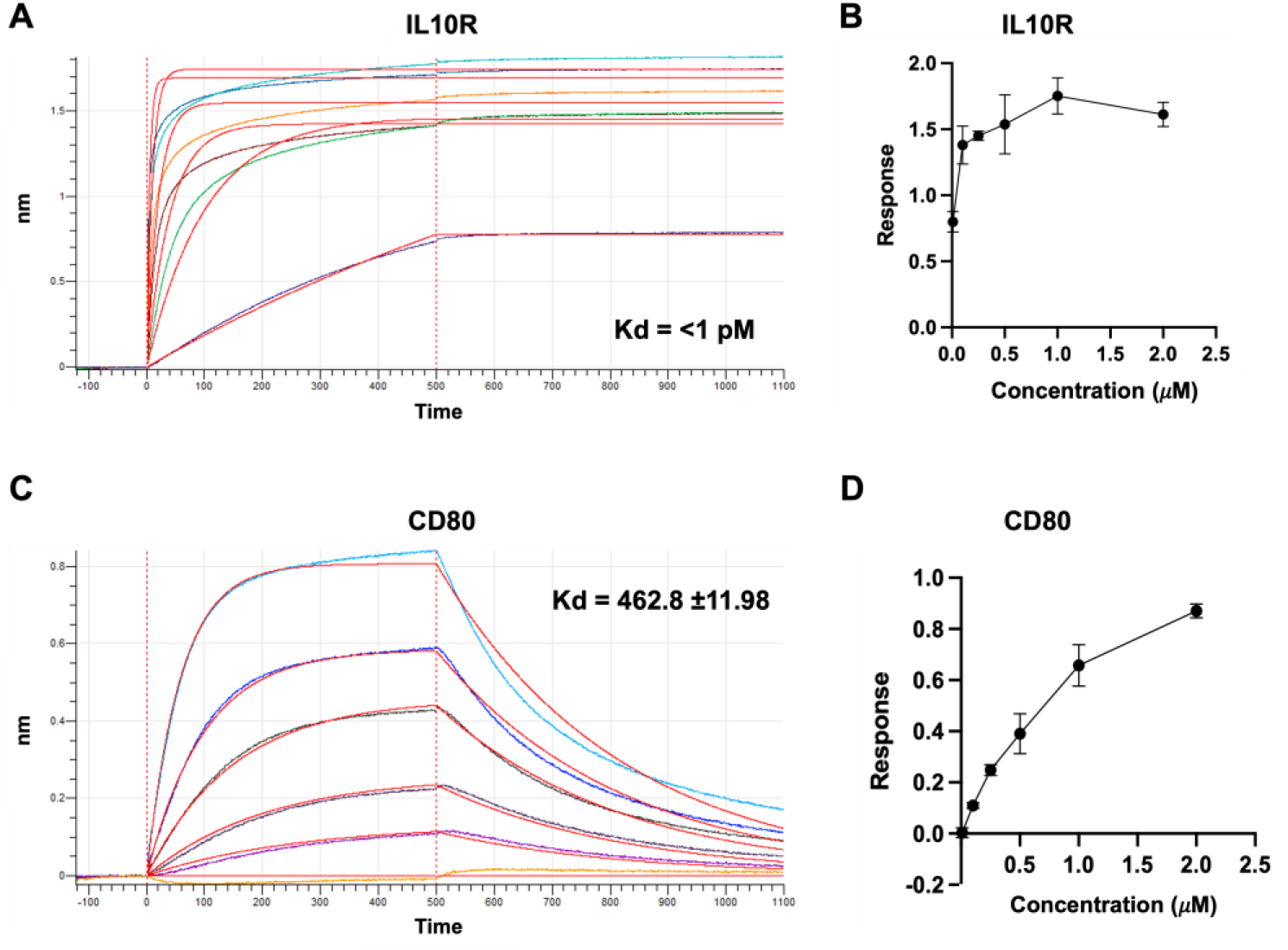
Kinetic binding analysis of the recombinant IL-10–IL-10–CTLA-4–Fc protein produced in mammalian cells by biolayer interferometry (BLI). **(A)** Representative sensorgrams and kinetic fits, and **(B)** steady-state binding analysis for binding to immobilized IL-10R. **(C)** Representative sensorgrams and kinetic fits, and **(D)** steady-state binding analysis for binding to immobilized CD80. Biotinylated receptors were loaded onto streptavidin biosensors and incubated with IL-10–IL-10–CTLA-4–Fc at concentrations ranging from 0.01 to 2 μM in PBS + 0.02% Tween-20. Data were globally fitted to a 1:1 binding model. Values represent mean ± SD from three independent experiments.

**Table 2.** Kinetic binding parameters for the mammalian-produced protein IL-10-IL-10/CTLA-4 interacting with immobilized CD80 or IL-10R measured by BLI. Values are mean ± SD from three independent experiments. KD is calculated as K_off_/K_on_.

| **IL10-IL10/CTLA4 produced in mammalian cells** | **Kd (nM)** | **Kon (1/Ms)** | **Koff (1/s)** |
| --- | --- | --- | --- |
| **Binding to CD80** | 462.8 ± 11.98 | 6.44E03 (±5.32E02) | 2.97E-03 (±0.0002) |
| **Binding to IL10R** | <1 pM | 9.52E04 (±6.9E03) | <1.0E-07 |

### 3.8 Evaluation of the effect of the chimera protein on IFNγ secretion of activated T-cells

In the final step, we evaluated the ability of virus-specific T cells (VSTs) to secrete IFNγ in response to the Adenovirus-derived Hexon antigen in the absence/presence of IL-10, CTLA-4, or the IL-10/CTLA-4 fusion protein. Here, we used APCs loaded with the Hexon peptide library at either 200 ng/mL (Figure 9A) or 1 ng/mL (Figure 9B) for antigen presentation by HLA class I and II molecules. After incubation for 2 hours, the APCs were washed of the antigens, and virus-specific T cells (VSTs) pre-incubated without or with the tested proteins at a final concentration of 200 nM were added to the APCs to form a co-culture. After 18 hours of incubation, the supernatant was collected and tested for IFNγ levels by ELISA (Fig. 9). Our results demonstrate that VSTs that were exposed to the Hexon antigen at 200 ng/ ml (9A) and 1 ng/ ml (9B) both activated the VSTs and secreted high levels of IFNγ as compared to the no antigen group, as expected. However, in the presence of IL-10, CTLA-4, or IL-10/CTLA-4 fusion drugs, VSTs activation was significantly inhibited, and IFNγ levels were significantly lower than in activated cells not treated with the proteins.

**Figure 9.**
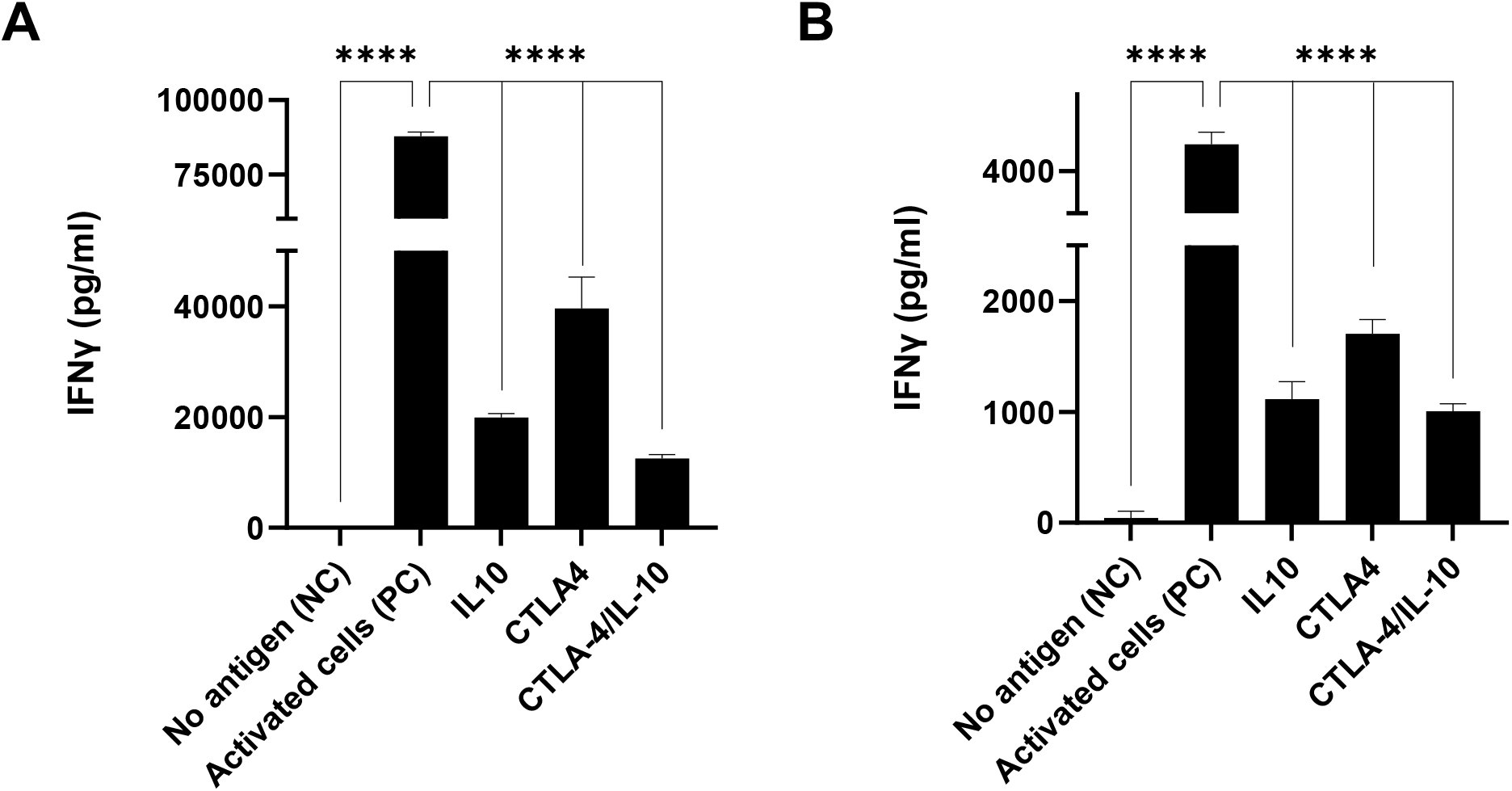
Effect of the CTLA-4/IL-10 protein on IFNγ secretion of virus-specific T cells. Co-culture of VST with APCs loaded with (A) 200 ng/ml or (B) 1 ng/ml of the Hexon peptide library, and were treated with IL-10, CTLA-4, or CTLA-4/IL-10 protein. Legend - No antigen (NC): VSTs that were not exposed to Hexon antigen; Activated cells (PC): VSTs that were exposed to the Hexon antigen with no addition of protein. IL-10, CTLA-4, or CTLA-4/IL-10: VSTs that were exposed to the Hexon antigen and the specific protein. Data are presented as mean ± SD. Statistical significance was determined by one-way ANOVA using GraphPad Prism (****, p < 0.0001).

## 4. Discussion

In this study, we engineered and characterized a bispecific CTLA-4/IL-10 fusion protein using both bacterial and mammalian expression systems. Beyond dual receptor engagement, the core biological hypothesis of the CTLA-4/IL-10 fusion is to create a localized immunoregulatory at CD80/86-expressing APCs. CTLA-4 is expected to tether the molecule to activated APCs while simultaneously antagonizing CD28-dependent stimulation, whereas IL-10 can trigger STAT3-driven anti-inflammatory programs in APCs and nearby immune cells. The mechanistic rationale for this dual-targeting approach is substantiated by prior studies on APVO210, a fusion of monomeric IL-10 and an anti-CD86 antibody [42]. While APVO210 demonstrated a promising activity profile, its clinical development was ultimately discontinued. Here, we propose that conjugating IL-10 to the CTLA-4 ECD offers an alternative to antibody-based design. Since CTLA-4-Fc is already a clinically validated scaffold, it provides a robust foundation for this design. Furthermore, this non-antibody architecture addresses key limitations of monoclonal antibodies (mAbs). Despite their dominance as pharmaceutical biologics, mAbs are hindered by high manufacturing costs, potential for immune-related adverse events, and limited tissue penetration due to their large molecular size[43, 44]. In contrast, engineered proteins based on receptor ECDs have demonstrated high binding affinity, superior tissue distribution [45-48], and a potentially safer immunogenicity profile [49].

Our study evaluated protein from bacterial and mammal sources. The bacterial platform provided a rapid, cost-effective route to functional protein, achieving a yield of ∼10 mg/L and demonstrating successful receptor recognition [50]. However, the lack of post-translational modifications (PTMs) in *E. coli* represents a significant limitation [51]. The inhibitory function of CTLA-4 is modulated by N-linked glycosylation, while the structural formation of IL-10 relies on disulfide bonds. Consequently, the mammalian CHO system offers an advantage by supporting these essential PTMs. Furthermore, the inclusion of an Fc-domain in the mammalian construct enhances stability and extends plasma half-life, a necessary feature for future pharmaceutical applications. Moreover, the CHO system delivered a superior yield of approximately 39 mg/L.

The binding results achieved with ELISA and BLI confirm that both receptor-binding domains in the chimera retain their structural integrity and biological activity following bacterial expression and refolding. The bacterial protein has a K_D_ of 155.3 nM towards CD80 and 45.7 nM towards IL-10R. Interestingly, the protein produced in mammalian cells has a KD of 462.8 nM towards CD80, but a much higher affinity of sub-nanomolar towards IL-10R. Whereas the strong affinity of the dimer IL-10 within the same polypeptide chain is expected, the protein produced in mammalian cells displayed slightly weaker binding to CD80 compared with the bacterially refolded version. Similar to IL10, CTLA4 also binds its receptors as a homodimer. As indicated by the analysis in Figure 6, dimerization of the protein occurs but it may be driven via FC-dimerization rather than spontaneous CTLA-4 interaction. Therefore, the difference in affinity could arise from steric constraints imposed by Fc dimerization and the tandem IL-10 configuration, which may alter the accessibility or orientation of the CTLA-4 ECD toward CD80. In addition, glycosylation heterogeneity on CTLA-4 could modulate association kinetics or introduce conformational ensembles that reduce apparent affinity under immobilized-receptor assay formats. An additional possibility is the selection of the linker for the chimera protein. Although glycine-serine linkers are commonly used due to their flexibility, it was demonstrated that altering the glycine-serine repetitions can impact the fusion protein’s stability and solubility [52, 53].

While ELISA and BLI demonstrate receptor binding, they do not establish functional immunomodulation. We further showed that the CTLA-4/IL-10 fusion protein successfully inhibits T-cell activation in a human VST model. To mimic natural human T cell function, we tested the VSTs against APCs loaded with moderate and low Hexon concentrations of 200 ng/ml and 1 ng/ml, respectively. Our results confirm that the protein not only binds its cognate receptors but also imparts functional cellular activity. An important next step is to verify downstream signaling, specifically CTLA-4–mediated inhibition and IL-10–dependent signaling events such as STAT3 phosphorylation. Additionally, a promising future direction for optimizing the CTLA-4/IL-10 bispecific protein involves rational protein engineering to tune the affinity of individual domains. Specifically, engineering a high-affinity CTLA-4 variant could enhance tissue localization by strongly anchoring the molecule to CD80/86-positive cells, thereby minimizing off-target interactions. Conversely, introducing mutations to attenuate IL-10 affinity [54] could mitigate the systemic toxicity or unpredictable co-stimulatory effects of wild-type IL-10. These design strategies warrant further investigation to advance the chimeric protein toward clinical therapeutic applications.

## 5. Conclusions

In this study, CTLA-4/IL-10 was expressed in bacterial and mammalian cells. Both protein sources bound the cognate receptors CD80 and IL-10R. The protein successfully inhibits T cell activation, as demonstrated by reducing IFNγ secretion levels in an in vitro T cell model of VSTs, showing its potential for cellular and therapeutic applications.

## Acknowledgments

This work was supported by funding from the Israeli Innovation Authority (IIA grant# 75654), the Israeli Ministry of Science, Technology and Innovation and Teva Pharmaceutical Ltd. Adi Cohen is grateful for the Ph.D. scholarship financial support from the Longevity and Health Research Center at Tel Aviv University. Matan Gabay is grateful to the Marian Gertner Institute for Medical Nano systems and the Yoran Institute for Advanced PhD Students in Personalized Medicine.

## Ethical approval

An IRB approval has been granted by the Sheba Medical Center Helsinki committee (7001-20-SMC) for the use of PBMCs and the establishment of virus-specific T cells.

## CRediT authorship contribution statement

**Adi Cohen:** Methodology, Formal analysis, Investigation, Data Curation, Validation, Writing original draft, Writing – Review and Editing; **Matan Gabay:** Methodology, Formal analysis, Investigation, Data Curation, Validation, Writing original draft; **Omri Cohen:** Methodology, Investigation; **Marina Sova:** Methodology, Resources, Project Administration; **Amihai Lieberman:** Methodology, Formal analysis, Investigation, Data Curation; Writing – Review and Editing; **Anat Shemer:** Methodology, Investigation; **Nira Varda-Bloom:** Methodology, Investigation; **Elad Jacoby:** Methodology, Investigation; **Gal Cafri:** Methodology, Validation, Supervision, Writing – Review and Editing; **Dorit Avni:** Methodology, Validation, Supervision, Funding acquisition, Writing – Review and Editing; **Itamar Yadid:** conceptualization, Validation, Supervision, Funding acquisition, Writing original draft, Writing – Review and Editing; **Maayan Gal:** conceptualization, Validation, Supervision, Funding acquisition, Writing original draft, Writing – Review and Editing;

