## Supplementary material for "Dual Immune Checkpoint and Cytokine Receptor Modulation by an Engineered Human CTLA-4/IL-10 Bispecific Fusion Protein": Supp

**Table S1. DNA sequence of GST-CTLA4-IL10-6xHis.**

| |
|---|
| ATGTCCCCTATACTAGGTTATTGGAAAATTAAGGGCCTTGTGCAACCCACTCGACTTCTTTTGGAATATCTT<br>GAAGAAAAATATGAAGAGCATTTGTATGAGCGCGATGAAGGTGATAAATGGCGAAACAAAAAGTTTGAATT<br>GGGTTTGGAGTTTCCCAATCTTCCTTATTATATTGATGGTGATGTTAAATTAACACAGTCTATGGCCATCAT<br>ACGTTATATAGCTGACAAGCACAACATGTTGGGTGGTTGTCCAAAAGAGCGTGCAGAGATTTCAATGCTT<br>GAAGGAGCGGTTTTGGATATTAGATACGGTGTTTCGAGAATTGCATATAGTAAAGACTTTGAAACTCTCAA<br>AGTTGATTTTCTTAGCAAGCTACCTGAAATGCTGAAAATGTTCGAAGATCGTTTATGTCATAAAACATATTT<br>AAATGGTGATCATGTAACCCATCCTGACTTCATGTTGTATGACGCTCTTGATGTTGTTTTATACATGGACCC<br>AATGTGCCTGGATGCGTTCCCAAAATTAGTTTGTTTTAAAAAACGTATTGAAGCTATCCCACAAATTGATAA<br>GTACTTGAAATCCAGCAAGTATATAGCATGGCCTTTGCAGGGCTGGCAAGCCACGTTTGGTGGTGGCGA<br>CCATCCTCCAAAATCGGATCTGGAAGTTCTGTTCCAGGGGCCCCTGGGATCGCATGTGGCTCAGCCTGC<br>CGTGGTCCTGGCCAGCTCCCGGGGCATCGCCTCTTTCGTGTGCGAGTACGCCAGCCCTGGCAAGGCAA<br>CCGAGGTGAGGGTGACAGTGCTGAGGCAGGCAGACAGCCAGGTGACCGAGGTGTGCGCAGCAACATAT<br>ATGATGGGCAACGAGCTGACCTTTCTGGACGATTCCATCTGTACCGGCACATCTAGCGGCAACCAAGTGA<br>ATCTGACCATCCAGGGCCTGAGAGCTATGGACACAGGCCTGTACATCTGTAAGGTGGAGCTGATGTATCC<br>CCCTCCATACTATCTGGGCATCGGCAACGGCACACAAATCTACGTGATCGATCCAGAGCCTTGCCCAGGA<br>GGAGGAGGCTCCTCTCCAGGCCAGGGCACCCAGAGCGAGAACAGCTGTACACACTTCCCCGGCAACCT<br>GCCTAATATGCTGCGGGACCTGAGAGATGCCTTTTCCAGGGTGAAGACCTTCTTTCAGATGAAGGACCAG<br>CTGGATAACCTGCTGCTGAAGGAGTCCCTGCTGGAGGACTTCAAGGGCTACCTGGGCTGCCAGGCCCTG<br>TCTGAGATGATCCAGTTTTATCTGGAGGAAGTGATGCCACAGGCCGAGAATCAGGACCCCGACATCAAG<br>GCCCACGTGAACTCTCTGGGCGAGAATCTGAAGACACTGAGGCTGAGGCTGCGGAGATGCCACCGGTTC<br>CTGCCCTGTGAGAACAAGTCTAAGGCCGTGGAGCAGGTGAAGAACGCCTTTAATAAGCTGCAGGAGAAG<br>GGCATCTATAAGGCCATGAGCGAGTTCGACATCTTCATCAATTACATTGAAGCCTACATGACTATGAAGAT<br>TAGAAACGGCAGCCACCATCACCATCACCATTAA |

**Table S2. Protein sequence of GST-CTLA4-IL10-6xHis.**

| |
|---|
| MSPILGYWKIKGLVQPTRLLLEYLEEKYEEHLYERDEGDKWRNKKFELGLEFPNLPYYIDGDVKLTQSMAIIR<br>YIADKHNMLGGCPKERAEISMLEGAVLDIRYGVSRIAYSKDFETLKVDFLSKLPEMLKMFEDRLCHKTYLNG<br>DHVTHPDFMLYDALDVVLYMDPMCLDAFPKLVCFKKRIEAIPQIDKYLKSSKYIAWPLQGWQATFGGGDHP<br>PKSDLEVLFQGPLGSHVAQPAVVLASSRGIASFVCEYASPGKATEVRVTVLRQADSQVTEVCAATYMMGN<br>ELTFLDDSICTGTSSGNQVNLTIQGLRAMDTGLYICKVELMYPPPYYLGIGNGTQIYVIDPEPCPGGGGSSP<br>GQGTQSENSCTHFPGNLPNMLRDLRDAFSRVKTFFQMKDQLDNLLLKESLLEDFKGYLGCQALSEMIQFYL<br>EEVMPQAENQDPDIKAHVNSLGENLKTLRLRLRRCHRFLPCENKSKAVEQVKNAFNKLQEKGIYKAMSEFD<br>IFINYIEAYMTMKIRNGSHHHHHH* |

Green- GST, blue- GRV 3C site, purple- CTLA4, orange- glycine-serine linker, red- IL10 and yellow- 6xHis tag.

**Table S3. DNA sequence of mammalian protein IL10-IL10-CTLA-4-Fc.**

GTTAGGCGTTTTGCGCTGCTTCGCGATGTACGGGCCAGATATACGCGTTGACATTGATTATTGACTAGTTA
TTAATAGTAATCAATTACGGGGTCATTAGTTCATAGCCCATATATGGAGTTCCGCGTTACATAACTTACGGT
AAATGGCCCGCCTGGCTGACCGCCCAACGACCCCCGCCCATTGACGTCAATAATGACGTATGTTCCCATA
GTAACGCCAATAGGGACTTTCCATTGACGTCAATGGGTGGAGTATTTACGGTAAACTGCCCACTTGGCAG
TACATCAAGTGTATCATATGCCAAGTACGCCCCCTATTGACGTCAATGACGGTAAATGGCCCGCCTGGCA
TTATGCCCAGTACATGACCTTATGGGACTTTCCTACTTGGCAGTACATCTACGTATTAGTCATCGCTATTAC
CATGGTGATGCGGTTTTGGCAGTACATCAATGGGCGTGGATAGCGGTTTGACTCACGGGGATTTCCAAGT
CTCCACCCCATTGACGTCAATGGGAGTTTGTTTTGGCACCAAAATCAACGGGACTTTCCAAAATGTCGTAA
CAACTCCGCCCCATTGACGCAAATGGGCGGTAGGCGTGTACGGTGGGAGGTCTATATAAGCAGAGCTCG
TTTAGTGAACCGTCAGATCGCCTGGAGACGCCATCCACGCTGTTTTGACCTCCATAGAAGACACCGGGAC
CGATCCAGCCTCCGGACTCTAGAGGATCGAACCCTTGAATTCCCGCCGCCACCATGGGTTGGAGTTGCA
TCATCCTATTTCTAGTGGCCACCGCTACCGGCGTGCACTCCGGCCCATCCCCTGGACAGGGCACCCAGT
CCGAGAACTCCTGTACCCACTTTCCTGGCAATCTGCCTAACATGCTGCGCGATCTGAGAGACGCCTTCTC
TAGAGTGAAGACCTTCTTCCAGATGAAGGACCAGCTGGACAACCTGCTGTTGAAAGAGAGCCTGCTGGAA
GATTTCAAGGGCTACCTGGGCTGTCAGGCCCTGTCCGAGATGATCCAGTTCTACCTGGAAGAGGTGATG
CCTCAGGCTGAGAACCAGGACCCTGACATCAAAGCTCATGTGAACTCTCTGGGCGAGAACCTGAAGACA
CTGAGACTGCGGCTCCGGCGGTGTCACAGATTCCTGCCCTGCGAGAACAAGTCTAAGGCCGTGGAACAG
GTGAAGAACGCCTTCAACAAGCTGCAAGAGAAAGGGATCTACAAGGCCATGTCCGAGTTCGACATCTTCA
TCAACTACATCGAGGCTTACATGACAATGAAGATCCGGAACGGCGGCGGCTCTGGTGGCGGCAGCCCTG
GCCAGGGAACACAGAGCGAGAACTCCTGCACACACTTCCCAGGCAATCTGCCTAACATGCTGAGAGACC
TGAGAGATGCCTTTTCTCGGGTGAAGACATTCTTCCAGATGAAGGATCAGCTGGATAACCTGCTGCTGAA
AGAGTCTCTCCTGGAGGACTTTAAGGGCTACCTCGGCTGCCAGGCTCTGTCCGAGATGATCCAGTTCTAC
CTGGAAGAAGTGATGCCTCAGGCCGAGAACCAGGATCCTGATATCAAGGCCCACGTGAACTCCCTGGGC
GAAAACCTGAAGACCCTGAGACTCCGGCTGAGACGGTGCCACCGGTTCCTGCCTTGTGAAAATAAGTCC
AAGGCCGTGGAACAAGTGAAAAACGCTTTTAACAAGCTACAAGAGAAGGGCATCTACAAGGCCATGTCTG
AGTTCGACATCTTTATCAACTATATTGAGGCCTACATGACCATGAAGATCAGAAACGGCGGCGGAGGCAG
TAGCGGCGGCGGTGGCTCCTCTTCCGAAAACCTGTACTTCCAAGGCCACGTGGCTCAGCCTGCTGTGGT
CCTGGCCTCTTCCAGAGGAATCGCCAGCTTTGTGTGCGAGTACGCCTCCCCAGGCAAGGCTACCGAGGT
CAGAGTCACCGTGCTGAGACAGGCCGACTCTCAGGTCACTGAAGTGTGCGCCGCTACGTACATGATGGG
CAACGAGCTGACCTTCCTGGACGACTCCATCTGCACCGGAACCTCTTCTGGCAACCAGGTGAACCTGAC
CATCCAGGGCCTGCGGGCTATGGACACCGGCCTGTACATCTGCAAAGTGGAACTGATGTATCCTCCTCC
CTACTACCTGGGAATCGGCAATGGCACACAGATCTACGTGATCGACCCCGAACCTTGTCCTGACTCCGAC
GGCGGCGGCTCCAGAGGCCCTACCATCAAGCCTTGCCCCCCTTGCAAGTGCCCTGCCCCCAACCTGCT
GGGAGGACCCTCTGTGTTCATCTTCCCTCCTAAGATCAAGGACGTGCTGATGATCAGCCTGTCTCCTATC
GTGACCTGTGTGGTCGTGGACGTGTCTGAGGATGACCCCGATGTGCAGATCTCTTGGTTCGTGAACAAC
GTGGAAGTCCACACAGCCCAGACCCAGACCCATAGAGAGGACTACAATTCTACCCTGAGAGTGGTGTCC
GCCCTGCCTATTCAGCACCAGGACTGGATGTCCGGAAAAGAGTTCAAGTGCAAAGTGAACAACAAGGAC
CTCCCAGCTCCTATCGAGCGCACCATCTCCAAGCCCAAGGGATCCGTGCGGGCCCCTCAAGTGTACGTG
CTGCCTCCACCTGAGGAAGAAATGACCAAGAAGCAGGTTACACTGACCTGCATGGTGACCGACTTCATGC
CCGAGGATATCTATGTGGAGTGGACCAACAACGGCAAGACCGAGCTGAACTACAAGAACACCGAGCCTG
TGCTGGACAGCGATGGCTCCTACTTCATGTACTCCAAGCTGAGGGTCGAAAAAAAGAACTGGGTCGAGA
GAAACTCCTACTCTTGCTCCGTGGTGCACGAGGGCCTGCACAACCACCACACCACCAAGTCCTTCTCCAG
AACCCCTGGCAAAGGCTCCTACCCCTACGACGTGCCCGACTACGCTCACCACCATCATCACCACTGATAA
GCTTAAGGGTTCGATCCCTACCGGTTAGTAATGAGTTTGATATCTCGACAATCAACCTCTGGATTACAAAA
TTTGTGAAAGATTGACTGGTATTCTTAACTATGTTGCTCCTTTTACGCTATGTGGATACGCTGCTTTAATGC
CTTTGTATCATGCTATTGCTTCCCGTATGGCTTTCATTTTCTCCTCCTTGTATAAATCCTGGTTGCTGTCTC
TTTATGAGGAGTTGTGGCCCGTTGTCAGGCAACGTGGCGTGGTGTGCACTGTGTTTGCTGACGCAACCC
CCACTGGTTGGGGCATTGCCACCACCTGTCAGCTCCTTTCCGGGACTTTCGCTTTCCCCCTCCCTATTGC
CACGGCGGAACTCATCGCCGCCTGCCTTGCCCGCTGCTGGACAGGGGCTCGGCTGTTGGGCACTGACA
ATTCCGTGGTGTTGTCGGGGAAGCTGACGTCCTTTCCATGGCTGCTCGCCTGTGTTGCCACCTGGATTCT
GCGCGGGACGTCCTTCTGCTACGTCCCTTCGGCCCTCAATCCAGCGGACCTTCCTTCCCGCGGCCTGCT
GCCGGCTCTGCGGCCTCTTCCGCGTCTTCGCCTTCGCCCTCAGACGAGTCGGATCTCCCTTTGGGCCGC
CTCCCCGCCTGGAACGGGGGAGGCTAACTGAAACACGGAAGGAGACAATACCGGAAGGAACCCGCGCT
ATGACGGCAATAAAAAGACAGAATAAAACGCACGGGTGTTGGGTCGTTTGTTCATAAACGCGGGGTTCGG
TCCCAGGGCTGGCACTCTGTCGATACCCCACCGAGACCCCATTGGGGCCAATACGCCCGCGTTTCTTCC
TTTTCCCCACCCCACCCCCCAAGTTCGGGTGAAGGCCCAGGGCTCGCAGCCAACGTCGGGGCGGCAGG
CCCTGCCATAGCAGATCTGCGCAGCTGGGGCTCTAGGGGGTATCCCCACGCGCCCTGTAGCGGCGCAT
TAAGCGCGGCGGGTGTGGTGGTTACGCGCAGCGTGACCGCTACACTTGCCAGCGCCCTAGCGCCCGCT
CCTTTCGCTTTCTTCCCTTCCTTTCTCGCCACGTTCGCCGGCTTTCCCCGTCAAGCTCTAAATCGGGGGC
TCCCTTTAGGGTTCCGATTTAGTGCTTTACGGCACCTCGACCCCAAAAAACTTGATTAGGGTGATGGTTCA
CGTAGTGGGCCATCGCCCTGATAGACGGTTTTTCGCCCTTTGACGTTGGAGTCCACGTTCTTTAATAGTG
GACTCTTGTTCCAAACTGGAACAACACTCAACCCTATCTCGGTCTATTCTTTTGATTTATAAGGGATTTTGC
CGATTTCGGCCTATTGGTTAAAAAATGAGCTGATTTAACAAAAATTTAACGCGAATTAATTCTGTGGAATGT

GTGTCAGTTAGGGTGTGGAAAGTCCCCAGGCTCCCCAGCAGGCAGAAGTATGCAAAGCATGCATCTCAA
TTAGTCAGCAACCAGGTGTGGAAAGTCCCCAGGCTCCCCAGCAGGCAGAAGTATGCAAAGCATGCATCT
CAATTAGTCAGCAACCATAGTCCCGCCCCTAACTCCGCCCATCCCGCCCCTAACTCCGCCCAGTTCCGCC
CATTCTCCGCCCCATGGCTGACTAATTTTTTTTATTTATGCAGAGGCCGAGGCCGCCTCTGCCTCTGAGCT
ATTCCAGAAGTAGTGAGGAGGCTTTTTTGGAGGCCTAGGCTTTTGCAAAAAGCTCCCGGGAGCTTGTATA
TCCATTTTCGGATCTGATCAAGAGACAGGATGAGGATCGTTTCGCATGATTGAACAAGATGGATTGCACG
CAGGTTCTCCGGCCGCTTGGGTGGAGAGGCTATTCGGCTATGACTGGGCACAACAGACAATCGGCTGCT
CTGATGCCGCCGTGTTCCGGCTGTCAGCGCAGGGGCGCCCGGTTCTTTTTGTCAAGACCGACCTGTCCG
GTGCCCTGAATGAACTGCAGGACGAGGCAGCGCGGCTATCGTGGCTGGCCACGACGGGCGTTCCTTGC
GCAGCTGTGCTCGACGTTGTCACTGAAGCGGGAAGGGACTGGCTGCTATTGGGCGAAGTGCCGGGGCA
GGATCTCCTGTCATCTCACCTTGCTCCTGCCGAGAAAGTATCCATCATGGCTGATGCAATGCGGCGGCTG
CATACGCTTGATCCGGCTACCTGCCCATTCGACCACCAAGCGAAACATCGCATCGAGCGAGCACGTACT
CGGATGGAAGCCGGTCTTGTCGATCAGGATGATCTGGACGAAGAGCATCAGGGGCTCGCGCCAGCCGA
ACTGTTCGCCAGGCTCAAGGCGCGCATGCCCGACGGCGAGGATCTCGTCGTGACCCATGGCGATGCCT
GCTTGCCGAATATCATGGTGGAAAATGGCCGCTTTTCTGGATTCATCGACTGTGGCCGGCTGGGTGTGG
CGGACCGCTATCAGGACATAGCGTTGGCTACCCGTGATATTGCTGAAGAGCTTGGCGGCGAATGGGCTG
ACCGCTTCCTCGTGCTTTACGGTATCGCCGCTCCCGATTCGCAGCGCATCGCCTTCTATCGCCTTCTTGA
CGAGTTCTTCTGAGCGGGACTCTGGGGTTCGCGAAATGACCGACCAAGCGACGCCCAACCTGCCATCAC
GAGATTTCGATTCCACCGCCGCCTTCTATGAAAGGTTGGGCTTCGGAATCGTTTTCCGGGACGCCGGCT
GGATGATCCTCCAGCGCGGGGATCTCATGCTGGAGTTCTTCGCCCACCCCAACTTGTTTATTGCAGCTTA
TAATGGTTACAAATAAAGCAATAGCATCACAAATTTCACAAATAAAGCATTTTTTTCACTGCATTCTAGTTGT
GGTTTGTCCAAACTCATCAATGTATCTTATCATGTCTGTATACCGTCGACCTCTAGCTAGAGCTTGGCGTA
ATCATGGTCATAGCTGTTTCCTGTGTGAAATTGTTATCCGCTCACAATTCCACACAACATACGAGCCGGAA
GCATAAAGTGTAAAGCCTGGGGTGCCTAATGAGTGAGCTAACTCACATTAATTGCGTTGCGCTCACTGCC
CGCTTTCCAGTCGGGAAACCTGTCGTGCCAGCTGCATTAATGAATCGGCCAACGCGCGGGGAGAGGCG
GTTTGCGTATTGGGCGCTCTTCCGCTTCCTCGCTCACTGACTCGCTGCGCTCGGTCGTTCGGCTGCGGC
GAGCGGTATCAGCTCACTCAAAGGCGGTAATACGGTTATCCACAGAATCAGGGGATAACGCAGGAAAGA
ACATGTGAGCAAAAGGCCAGCAAAAGGCCAGGAACCGTAAAAAGGCCGCGTTGCTGGCGTTTTTCCATA
GGCTCCGCCCCCCTGACGAGCATCACAAAAATCGACGCTCAAGTCAGAGGTGGCGAAACCCGACAGGAC
TATAAAGATACCAGGCGTTTCCCCCTGGAAGCTCCCTCGTGCGCTCTCCTGTTCCGACCCTGCCGCTTAC
CGGATACCTGTCCGCCTTTCTCCCTTCGGGAAGCGTGGCGCTTTCTCATAGCTCACGCTGTAGGTATCTC
AGTTCGGTGTAGGTCGTTCGCTCCAAGCTGGGCTGTGTGCACGAACCCCCCGTTCAGCCCGACCGCTGC
GCCTTATCCGGTAACTATCGTCTTGAGTCCAACCCGGTAAGACACGACTTATCGCCACTGGCAGCAGCCA
CTGGTAACAGGATTAGCAGAGCGAGGTATGTAGGCGGTGCTACAGAGTTCTTGAAGTGGTGGCCTAACT
ACGGCTACACTAGAAGAACAGTATTTGGTATCTGCGCTCTGCTGAAGCCAGTTACCTTCGGAAAAAGAGT
TGGTAGCTCTTGATCCGGCAAACAAACCACCGCTGGTAGCGGTGGTTTTTTTGTTTGCAAGCAGCAGATT
ACGCGCAGAAAAAAAGGATCTCAAGAAGATCCTTTGATCTTTTCTACGGGGTCTGACGCTCAGTGGAACG
AAAACTCACGTTAAGGGATTTTGGTCATGAGATTATCAAAAAGGATCTTCACCTAGATCCTTTTAAATTAAA
AATGAAGTTTTAAATCAATCTAAAGTATATATGAGTAAACTTGGTCTGACAGTTACCAATGCTTAATCAGTG
AGGCACCTATCTCAGCGATCTGTCTATTTCGTTCATCCATAGTTGCCTGACTCCCCGTCGTGTAGATAACT
ACGATACGGGAGGGCTTACCATCTGGCCCCAGTGCTGCAATGATACCGCGAGACCCACGCTCACCGGCT
CCAGATTTATCAGCAATAAACCAGCCAGCCGGAAGGGCCGAGCGCAGAAGTGGTCCTGCAACTTTATCC
GCCTCCATCCAGTCTATTAATTGTTGCCGGGAAGCTAGAGTAAGTAGTTCGCCAGTTAATAGTTTGCGCAA
CGTTGTTGCCATTGCTACAGGCATCGTGGTGTCACGCTCGTCGTTTGGTATGGCTTCATTCAGCTCCGGT
TCCCAACGATCAAGGCGAGTTACATGATCCCCCATGTTGTGCAAAAAAGCGGTTAGCTCCTTCGGTCCTC
CGATCGTTGTCAGAAGTAAGTTGGCCGCAGTGTTATCACTCATGGTTATGGCAGCACTGCATAATTCTCTT
ACTGTCATGCCATCCGTAAGATGCTTTTCTGTGACTGGTGAGTACTCAACCAAGTCATTCTGAGAATAGTG
TATGCGGCGACCGAGTTGCTCTTGCCCGGCGTCAATACGGGATAATACCGCGCCACATAGCAGAACTTTA
AAAGTGCTCATCATTGGAAAACGTTCTTCGGGGCGAAAACTCTCAAGGATCTTACCGCTGTTGAGATCCA
GTTCGATGTAACCCACTCGTGCACCCAACTGATCTTCAGCATCTTTTACTTTCACCAGCGTTTCTGGGTGA
GCAAAAACAGGAAGGCAAAATGCCGCAAAAAAGGGAATAAGGGCGACACGGAAATGTTGAATACTCATAC
TCTTCCTTTTTCAATATTATTGAAGCATTTATCAGGGTTATTGTCTCATGAGCGGATACATATTTGAATGTAT
TTAGAAAAATAAACAAATAGGGGTTCCGCGCACATTTCCCCGAAAAGTGCCACCTGACGTCGACGGATCG
GGAGATCTCCCGATCCCCTATGGTGCACTCTCAGTACAATCTGCTCTGATGCCGCATAGTTAAGCCAGTA
TCTGCTCCCTGCTTGTGTGTTGGAGGTCGCTGAGTAGTGCGCGAGCAAAATTTAAGCTACAACAAGGCAA
GGCTTGACCGACAATTGCATGAAGAATCTGCTTAGG

**Table S4. Protein sequence of mammalian protein IL10-IL10-CTLA4-Fc**

| |
|---|
| MGWSCIILFLVATATGVHSGPSPGQGTQSENSCTHFPGNLPNMLRDLRDAFSRVKTFFQMKDQLD<br>NLLLKESLLEDFKGYLGCQALSEMIQFYLEEVMPQAENQDPDIKAHVNSLGENLKTLRLRLRRCHR<br>FLPCENKSKAVEQVKNAFNKLQEKGIYKAMSEFDIFINYIEAYMTMKIRNGGGSGGGSPGQGTQSE<br>NSCTHFPGNLPNMLRDLRDAFSRVKTFFQMKDQLDNLLLKESLLEDFKGYLGCQALSEMIQFYLEE<br>VMPQAENQDPDIKAHVNSLGENLKTLRLRLRRCHRFLPCENKSKAVEQVKNAFNKLQEKGIYKAM<br>SEFDIFINYIEAYMTMKIRNGGGGSSGGGGSSSENLYFQGHVAQPAVVLASSRGIASFVCEYASPGK<br>ATEVRVTVLRQADSQVTEVCAATYMMGNELTFLDDSICTGTSSGNQVNLTIQGLRAMDTGLYICKVE<br>LMYPPPYYLGIGNGTQIYVIDPEPCPDSDGGGS**RGPTIKPCPPCKCPAPNLLGGPSVFIFPPKIKD**<br>**VLMISLSPIVTCVVVDVSEDDPDVQISWFVNNVEVHTAQTQTHREDYNSTLRVVSALPIQHQD**<br>**WMSGKEFKCKVNNKDLPAPIERTISKPKGSVRAPQVYVLPPPEEEMTKKQVTLTCMVTDFMPE**<br>**DIYVEWTNNGKTELNYKNTEPVLDSDGSYFMYSKLRVEKKNWVERNSYSCSVVHEGLHNHHT**<br>**TKSFSRTPGK**GSYPYDVPDYAHHHHHH |

Green- Secretion signal, purple- IL-10, Yellow – Linkers, Blue - CTLA4, Brown - FC, Red – Linker and Hisx6 tag.

**Table S5. Buffers used for bacterial protein purification and refolding**

| Buffer Name | Composition |
|---|---|
| Lysis Buffer | 50 mM Tris-HCl (pH 7.5), 500 mM NaCl, 10 mM EDTA, 5 mM DTT, 2% Triton X-100, protease inhibitor (1:500) |
| Denaturation Buffer | 20 mM Tris-HCl (pH 8.0), 0.5 M NaCl, 6 M guanidine hydrochloride |
| Equilibration Buffer | 20 mM Tris-HCl (pH 8.0), 500 mM NaCl, 5 mM imidazole, 6 M guanidine hydrochloride |
| Wash Buffer | 20 mM Tris-HCl (pH 8.0), 500 mM NaCl, 20 mM imidazole, 6 M urea |
| Elution Buffer | 20 mM Tris-HCl (pH 8.0), 500 mM NaCl, 6 M urea |
| Refolding Buffer | 55 mM Tris-HCl (pH 8.0), 880 mM L-arginine, 10 mM NaCl, 0.88 mM KCl, 1 mM EDTA, 1 mM cystine, 5 mM cysteine |
| Storage Buffer | Refolding Buffer supplemented with 20% (v/v) glycerol |

**Table S6. Primary and secondary antibodies used for ELISA and/or western blot.**

| Antibody | Function |
|---|---|
| Anti-6xHis tag (Abcam, ab18184) | Primary Ab in western blot |
| Anti-IL10 antibody (Abcam, ab134742) | Primary Ab in western blot |
| THE™ GST Antibody (A2S, A00865) | Primary Ab in western blot |
| Anti-CTLA4 antibody (Abcam, ab231949) | Primary Ab in western blot |
| Goat Anti-Rabbit IgG Antibody, HRP-conjugate (Merck,12-348 ) | Secondary Ab in western blot |
| Goat anti-mouse IgG Antibody, HRP conjugate (Merck, 12-349) | Secondary Ab in western blot |
| Goat Anti-Human IgG Fc-HRP (Abcam, ab98624) | Detection of Fc-IL10 in ELISA |
| Streptavidin-HRP (Abcam, ab7403) | Detection of biotinylated CD80 in ELISA |
